# PARP inhibition causes premature loss of cohesion in cancer cells

**DOI:** 10.1101/153932

**Authors:** Eva Kukolj, Tanja Kaufmann, Amalie E. Dick, Robert Zeillinger, Daniel W. Gerlich, Dea Slade

## Abstract

Poly(ADP-ribose) polymerases (PARPs) regulate various aspects of cellular function including mitotic progression. Although PARP inhibitors have been undergoing various clinical trials and the PARP1/2 inhibitor olaparib was approved as monotherapy for BRCA-mutated ovarian cancer, their mode of action in killing tumour cells is not fully understood. We investigated the effect of PARP inhibition on mitosis in cancerous (cervical, ovary, breast and osteosarcoma) and non-cancerous cells by live-cell imaging. The clinically relevant inhibitor olaparib induced strong perturbations in mitosis, including problems with chromosome alignment at the metaphase plate, anaphase delay, and premature loss of cohesion (cohesion fatigue) after a prolonged metaphase arrest, resulting in sister chromatid scattering. PARP1 and PARP2 depletion suppressed the phenotype while PARP2 overexpression enhanced it, suggesting that olaparib-bound PARP1 and PARP2 rather than the lack of catalytic activity causes this phenotype. Olaparib-induced mitotic chromatid scattering was observed in various cancer cell lines with increased protein levels of PARP1 and PARP2, but not in non-cancer or cancer cell lines that expressed lower levels of PARP1 or PARP2. Interestingly, the sister chromatid scattering phenotype occurred only when olaparib was added during the S-phase preceding mitosis, suggesting that PARP1 and PARP2 entrapment at replication forks impairs sister chromatid cohesion. Clinically relevant DNA-damaging agents that impair replication progression such as topoisomerase inhibitors and cisplatin were also found to induce sister chromatid scattering and metaphase plate alignment problems, suggesting that these mitotic phenotypes are a common outcome of replication perturbation.

## INTRODUCTION

Poly(ADP-ribose) polymerases (PARPs) are enzymes important for diverse cellular processes ranging from transcriptional regulation and cell-cycle control, to chromatin dynamics, DNA repair, mitosis and cell death [1-3]. PARPs synthesize poly(ADP-ribose) (PAR) from NAD by attaching ADP-ribose units via glycosidic ribose-ribose bonds onto themselves (auto-modification) and other protein acceptors (hetero-modification) [4]. PAR—a short-lived post-translational modification—is an ideal mediator of dynamic cellular processes based on the formation of interaction scaffolds or the disruption of protein-protein and protein-DNA interactions [5]. More than 90% of cellular PAR is synthesized by PARP1 as the most abundant and the most highly active PARP [6].

Regulation of various DNA repair pathways such as single-strand break repair (SSBR), homologous recombination (HR) and non-homologous end joining remains the best studied role of PARP1 [7, 8]. Additionally, PARP1 promotes replication fork reversal and HR-dependent restart of stalled or collapsed replication forks [9-12]. PARP inhibition causes stalling or collapse of replication forks, resulting in lethal double-strand DNA breaks (DSBs) [13]. Replication blockage is presumably a consequence of entrapment and accumulation of inactive PARP1 on DNA by NAD-mimicking PARP inhibitors [13]. Sensitization of HR-deficient cancer cells by PARP inhibition has given rise to synthetic lethality approaches, whereby pharmacological inhibition of one DNA repair pathway coupled with genetic defects in another pathway causes lethality due to inability to repair damaged DNA [14]. In the first example of synthetic lethality induced by PARP inhibition, PARP1/2 inhibitor was shown to induce chromosomal instability, cell cycle arrest and apoptosis in breast cancer patients carrying heterozygous loss-of-function *BRCA* mutations [15, 16]. Another example of synthetic lethality between PARP1 inhibition and cohesin mutations further corroborates the importance of PARP1 for replication fork stability [17].

In addition to DNA repair, the roles of PARPs in the regulation of inflammatory mediators, cellular energetics, cell fate, gene transcription, ERK-mediated signalling and mitosis might underlie the susceptibility of cancer cells to PARP inhibition [18]. PARPs have distinct mitotic functions. PARP1 and PARP2 localize at centromeres and interact with centromeric proteins [19]. PARP1 is required for the maintenance of the spindle assembly checkpoint and post-mitotic checkpoint; its depletion or inhibition result in centrosome amplification and aneuploidy [20-22]. PARP1 knock-out mouse oocytes exhibit incomplete synapsis of homologous chromosomes, deficient sister chromatid cohesion during metaphase II and failure to maintain metaphase arrest due to lack of centromeric recruitment of the mitotic checkpoint protein BUB3 [23]. The E3 ubiquitin ligase CHFR (checkpoint with FHA and RING finger domains) regulates the mitotic checkpoint via PARP1 ubiquitination and degradation during mitotic stress, resulting in cell cycle arrest in prophase [24]. Tankyrase (PARP5) has also been implicated in mitotic regulation; it is found around the pericentriolar matrix of mitotic chromosomes and was shown to regulate spindle assembly [25, 26] together with PARP3 [27].

Olaparib is the only PARP1/2 inhibitor approved for treatment of pretreated or platinum sensitive ovarian cancer associated with defective BRCA1/2 genes. Talazoparib is the most potent PARP1/2 inhibitor developed to date, exerting its cytotoxicity by PARP trapping rather than catalytic inhibition [28]. The catalytic inhibitory effect of talazoparib is comparable to olaparib; nevertheless, it is 100-fold more potent at trapping PARP-DNA complexes [28]. Veliparib is among the least potent PARP1/2 inhibitors with weak catalytic inhibition and low PARP trapping efficiency [13]. All three inhibitors are currently undergoing various clinical trials.

Considering the multiple roles of PARP in mitosis, we investigated the effect of PARP inhibition on mitotic progression by live-cell imaging. PARP1/2 inhibition with olaparib, talazoparib or veliparib induced metaphase arrest and sister chromatid scattering in HeLa cells, leading to cell death. Chromatid scattering in mitosis was caused by premature loss of cohesion in interphase cells whereby olaparib treatment caused a two-fold increase in sister chromatid distance. Premature loss of cohesion occurred when olaparib was added already during S-phase, suggesting that replication fork blockage due to PARP entrapment leads to loss of cohesion and subsequent defects in mitosis. Premature loss of cohesion was also observed in cancer cell lines of cervical, ovary, breast and osteosarcoma origin that exhibit S-phase stalling upon olaparib treatment. The severity of this mitotic phenotype across different cell lines correlated with PARP1 and PARP2 protein levels, was rescued by PARP1 or PARP2 depletion and exacerbated by PARP2 overexpression. Similar mitotic phenotypes were also found upon treatment with DNA-damaging agents that cause S-phase stalling such as topoisomerase inhibitors (camptothecin, etoposide) and cisplatin, suggesting that death by mitotic failure is a general phenomenon of perturbed replication.

## RESULTS

### Olaparib causes anaphase delay and chromatid scattering in metaphase-arrested cells

In order to investigate the effect of PARP inhibition on mitosis, we performed live-cell imaging of HeLa cells stably expressing H2B-mCherry together with securin-EGFP [29] treated with olaparib (AZD2281, Ku-0059436) [30], talazoparib (BMN 673) [31] or veliparib (ABT-888) [32] as PARP1/2 inhibitors, XAV-939 as tankyrase1/2 (PARP5a/b) inhibitor [33] and ME328 as PARP3 inhibitor (Figure 1A) [34]. Of the five tested inhibitors applied at different concentrations for 30 h, only PARP1/2 inhibitors caused anaphase delay measured as the time required for the cells to progress from nuclear envelope breakdown (NEBD) to anaphase (Figure 1A,B). The median NEBD-anaphase duration was extended from 42 min in DMSO-treated control cells to 57 min for 10 μM olaparib-treated cells, to 60 min for 30 μM veliparib and to 60 min for 100 nM talazoparib (Figure 1A,B). 40- 50% of inhibitor-treated mitotic cells failed to enter anaphase due to metaphase plate formation problems or chromatid scattering after correct metaphase plate formation (Figure 1B-D).

**Figure 1:**
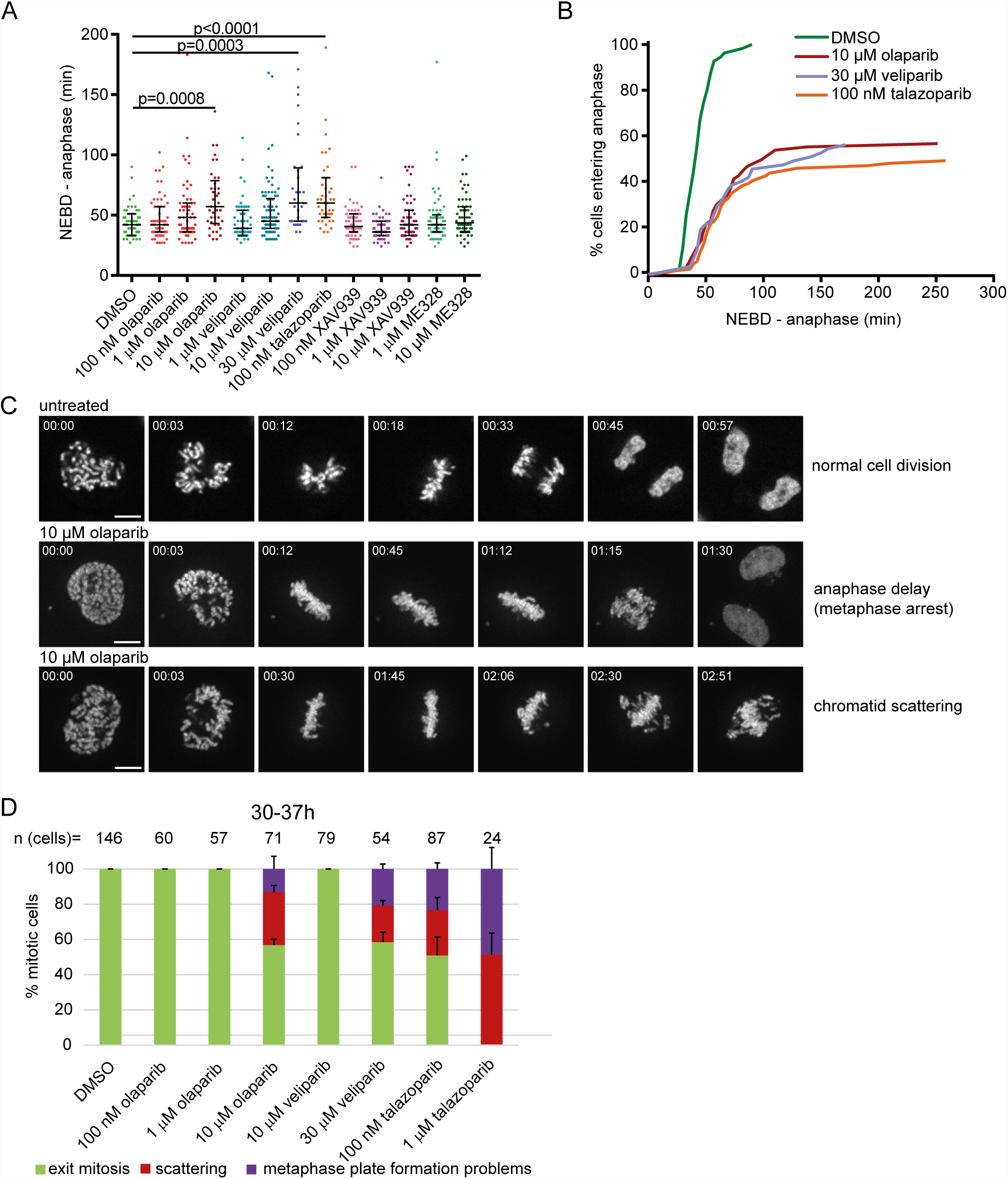
PARP1/2 inhibition causes anaphase delay and chromatid scattering (premature loss of cohesion) in HeLa cells. **(A)** Duration of NEBD-anaphase was analysed by live imaging of HeLa cells stably expressing H2B-mCherry EGFP-securin treated with PARP inhibitors for 30 h. Each dot represents one cell. **(B)** Cumulative frequency distribution representing both anaphase delay and inability to enter anaphase in cells treated with indicated concentrations of PARP inhibitors. **(C)** Stills of representative mitotic phenotypes. Scale bar= 10 μm. **(D)** Percentage of different mitotic phenotypes observed after treatment with PARP inhibitors.

Chromatid scattering was initially described as a result of prolonged metaphase in cells treated with the proteasome inhibitor MG132 or depleted of the anaphase promoting complex/cyclosome (APC/C) activator Cdc20, and thought to result from premature loss of cohesion (cohesion fatigue) [35, 36]. Chromatid scattering was observed in 30±4% of cells treated with 10 μM olaparib for 30-37 h, 21±3% of cells treated with 30 μM veliparib and 26±7% of cells treated with 100 nM talazoparib (Figure 1D). These cells were arrested in a metaphase-like state as revealed by high levels of cyclin B and securin (Supplementary Figure 1). Unless cyclin B and securin are ubiquitinated by the APC, separase-mediated cleavage of centromeric cohesion cannot promote metaphase to anaphase transition [37]. XAV-939 or ME328 did not induce the scattering phenotype.

### Chromatid scattering correlates with the efficiency of inhibition and S-phase stalling induced by PARP inhibitors

We next examined whether chromatid scattering is linked with the efficiency of PARP1 inhibition and its effect on cell cycle progression (Supplementary Figure 2). 10 μM olaparib, 30 μM veliparib and 100 nM olaparib efficiently inhibited PARP1 auto-PARylation as determined by Western blotting with an anti-PAR antibody (>90% reduction after 24 h compared to the DMSO-treated cells) (Supplementary Figure 2A). Although *in vitro* assays determined 1.4 nM and 12.3 nM as IC50 values of olaparib towards PARP1 and PARP2 [38], we found that olaparib has to be used at a much higher concentration in order to inhibit PARP1 in HeLa cells (Supplementary Figure 2A). However, the concentration of 10 μM olaparib that showed inhibition of PARP1 activity, anaphase delay and scattering is still significantly below the concentration of olaparib used in clinical trials (peak plasma concentration=24 μM) [39]. Our data confirm that talazoparib (IC_50_ values of 1.1 and 4.1 nM for PARP1 and PARP2, respectively) is a more potent inhibitor than olaparib [28, 38], whereas veliparib (IC_50_ values of 3.3 and 17.5 nM for PARP1 and PARP2, respectively) is a weaker inhibitor [13, 38]. Flow cytometry analysis confirmed that ≥1 μM olaparib, ≥100 nM talazoparib and ≥30 μM veliparib cause G2/M arrest [15, 40] (Supplementary Figure 2B). Depending on the incubation time, olaparib caused either stalling in S-phase (12 h incubation) or G2/M arrest (28 and 32 h of incubation) (Supplementary Figure 2C). Neither XAV-939 nor ME328 had an effect on cell cycle progression (Supplementary Figure 2B).

These results indicate that the concentrations of olaparib, talazoparib and veliparib that induce chromatid scattering strongly inhibit PARP1 activity and perturb S-phase progression. The strong S-phase stalling is presumably due to the trapping of PARP1 and PARP2 on DNA by these inhibitors [13, 28]. The inhibitors differ in their efficiency of catalytic inhibition, S-phase stalling and chromatid scattering, with talazoparib>olaparib>veliparib. Taken together, our data reveal that in addition to S-phase stalling and anaphase delay, prolonged olaparib, talazoparib or veliparib treatment induce chromatid scattering in HeLa cells.

### PARP silencing does not cause mitotic phenotypes found after PARP inhibition

PARP inhibition and depletion of PARP protein does not always yield the same phenotypes; PARP inhibition is more effective in inducing apoptosis in BRCA-deficient cells compared to PARP knock-down [16]; wild-type cells treated with olaparib are more sensitive to MMS than PARP1 knock-out cells [ 13]; PARP inhibition impairs SSB repair to a greater extent than PARP depletion [41, 42]; the level of DSBs is higher in olaparib-treated than PARP-depleted cells under basal conditions [13]. PARP inhibition was shown to cause S-phase stalling whereas PARP silencing had no effect [13]. To test whether depleting PARP causes the same mitotic phenotypes as PARP inhibition, we performed live-cell imaging 24 h after depleting individual PARP members by three different siRNAs (Supplementary Figure 3 and Supplementary Table 1). Anaphase delay was observed for only two siRNAs targeting PARP1, one siRNA targeting PARP2 and two siRNAs targeting PARP3 (Supplementary Figure 3A-C) (siControl: 33 min; siPARP1: 42 min, 39 min; siPARP2: 57 min; siPARP3: 58 min, 54 min), which suggests that it is an off-target effect. Silencing tankyrase 1 did not cause anaphase delay (Supplementary Figure 3D). None of the siRNAs caused scattering. This suggests that entrapment of inactive PARP1/2 at replication forks is likely to be the cause of mitotic phenotypes observed after PARP1/2 inhibition.

### Chromatid scattering is caused by specific inhibition of PARP1 and PARP2 by olaparib

PARP inhibition was previously shown to be more effective than PARP depletion and the cytotoxic effects of olaparib were directly linked with PARP1 rather than off-targets [13]. We also tested whether the scattering phenotype caused by PARP inhibition—but not depletion—is a specific or off-target effect of PARP inhibition (Figure 2). PARP1 depletion by RNAi reduced the degree of olaparib-induced scattering from 24±8 to 8±5% (Figure 2A). Both PARP2 and combined PARP1/2 depletion completely rescued olaparib-induced scattering (Figure 2A). This indicates that the scattering phenotype is caused by olaparib-inhibited PARP1 and PARP2, rather than an off-target inhibition by olaparib. Flow cytometry analysis additionally showed that PARP1 depletion in olaparib-treated cells reversed S-phase stalling, while PARP2 depletion had a partial effect (Figure 2B). Neither PARP1 nor PARP2 depletion rescued anaphase delay (Figure 2C). Importantly, concomitant PARP1 and PARP2 depletion increased survival of HeLa cells exposed to 10 μM olaparib for 72 h from 83±2 to 94±1% according to the MTS assay, which measures the percentage of metabolically active cells (Figure 2D).

**Figure 2:**
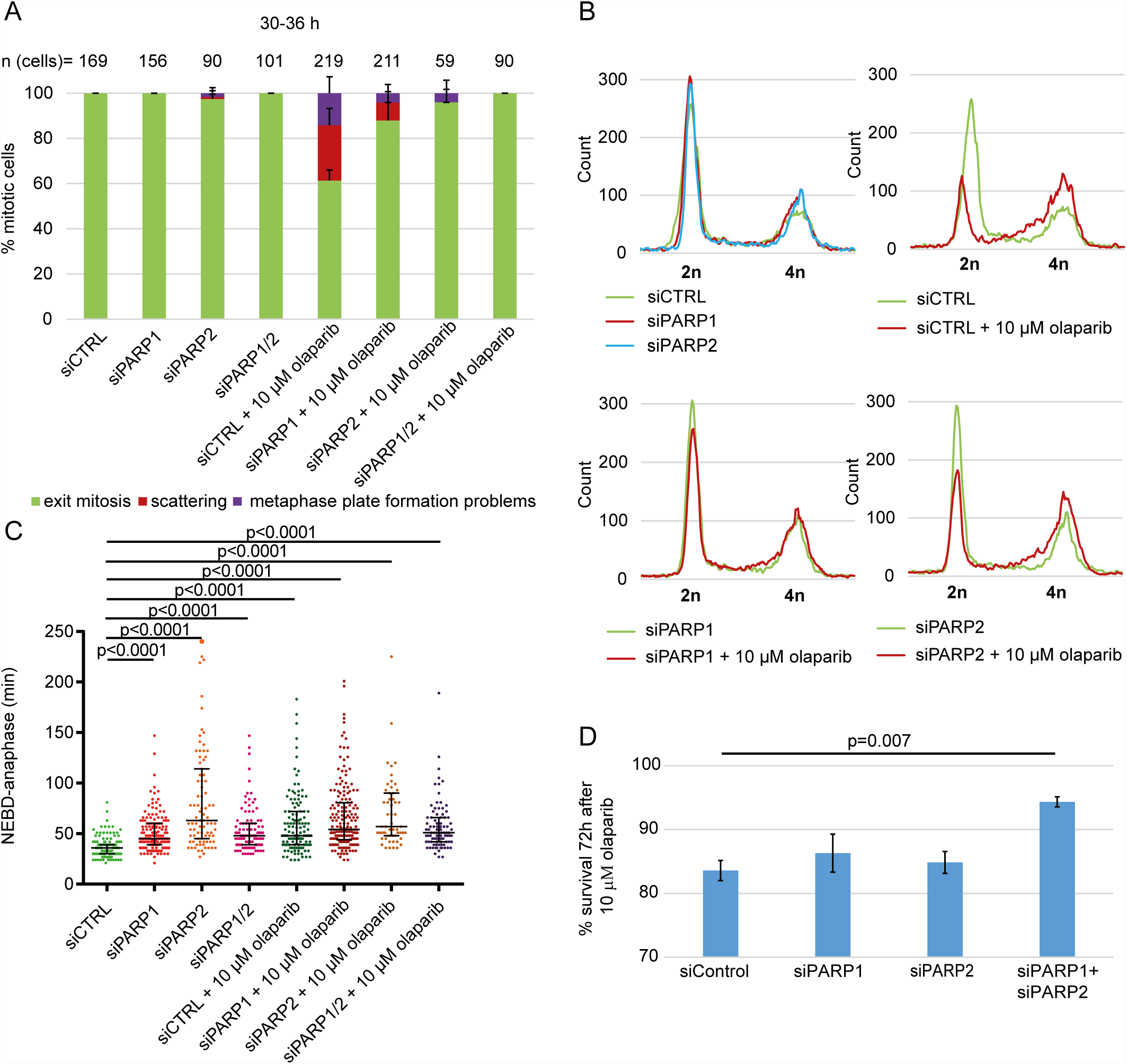
PARP1 and PARP2 depletion rescue olaparib-induced scattering, S-phase stalling and cytotoxicity but not anaphase delay in HeLa. **(A)** Percentage of different mitotic phenotypes, **(B)** flow cytometry analysis, **(C)** NEBD-anaphase duration and **(D)** cell survival after concomitant treatment with olaparib and siPARP1/2 compared to single treatments. siPARP1 E and siPARP2 E were used. Cell survival was determined using MTS assay.

Given that PARP1/2 silencing rescued olaparib-induced scattering, we also tested whether PARP1/2 overexpression can exacerbate the phenotype (Figure 3). We generated EGFP-PARP1 or EGFP-PARP2 overexpressing HeLa cells and performed live-cell imaging. PARP1 overexpression had no effect on scattering but slightly increased olaparib-induced anaphase delay; PARP2 increased the degree of scattering from 34±4% to 47±6% (p=0.037), without affecting anaphase delay (Figure 3). Taken together, our data indicate that (i) chromatid scattering is a specific outcome of olaparib-mediated PARP1 and PARP2 inhibition, (ii) PARP2 inhibition has a stronger effect on scattering, and (iii) olaparib cytotoxicity can be partly rescued by depleting PARP1 and PARP2.

**Figure 3:**
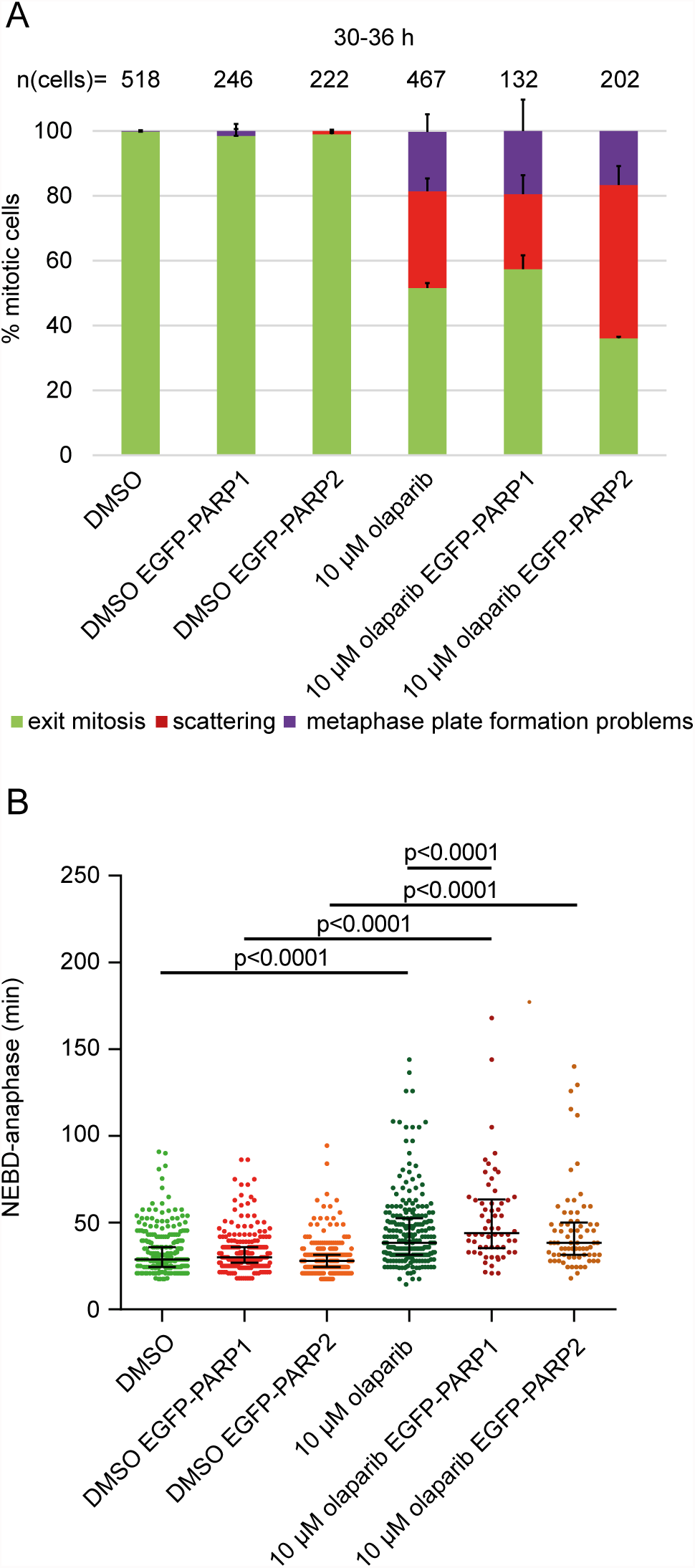
PARP2 overexpression exacerbates olaparib-induced scattering. **(A)** Percentage of different mitotic phenotypes and **(B)** NEBD-anaphase duration in EGFP-PARP1 or EGFP-PARP2-overexpressing HeLa cells imaged 30-36 h after treatment with 10 μM olaparib.

### Chromatid scattering is caused by the S-phase effects of olaparib

Olaparib did not perturb mitotic progression if added just before the onset of mitosis (Figure 4A), suggesting that defects from earlier cell cycle stages gave rise to the observed mitotic phenotypes. To test this, H2B-mCherry HeLa cells transfected with EGFP-PCNA were synchronized by double thymidine block (Figure 4). Olaparib was added (i) immediately after the release into S-phase and washed out after 6 h (estimated S-phase duration in HeLa cells according to EGFP-PCNA foci); (ii) immediately after the release into S-phase and washed out before mitotic entry; (iii) 30 min before mitosis (NEBD considered as mitotic entry) (Figure 4B). Olaparib was efficiently washed out as judged by the restoration of PARP1 auto-PARylation within two hours of olaparib removal (Figure 4C). Live-cell imaging on a spinning disc setup enabled us to distinguish with confidence between the sister chromatid scattering phenotype and the metaphase plate formation problems (Figure 4D). The presence of olaparib only during S-phase was sufficient to induce chromatid scattering and metaphase plate formation problems in 21±8% and 22±5% of mitotic cells, respectively (Figure 4D-E). Both phenotypes resulted in cell death (Figure 4D). Apoptotic death was observed in additional 11% of the analysed mitotic population as well as in 8% of untreated cells, and may be due to anaphase-related segregation defects, thymidine-induced damage or physical causes (e.g., phototoxicity during imaging) (Figure 4E). PARP inhibition during S-phase thus causes subsequent perturbations during mitosis leading to cell death.

**Figure 4:**
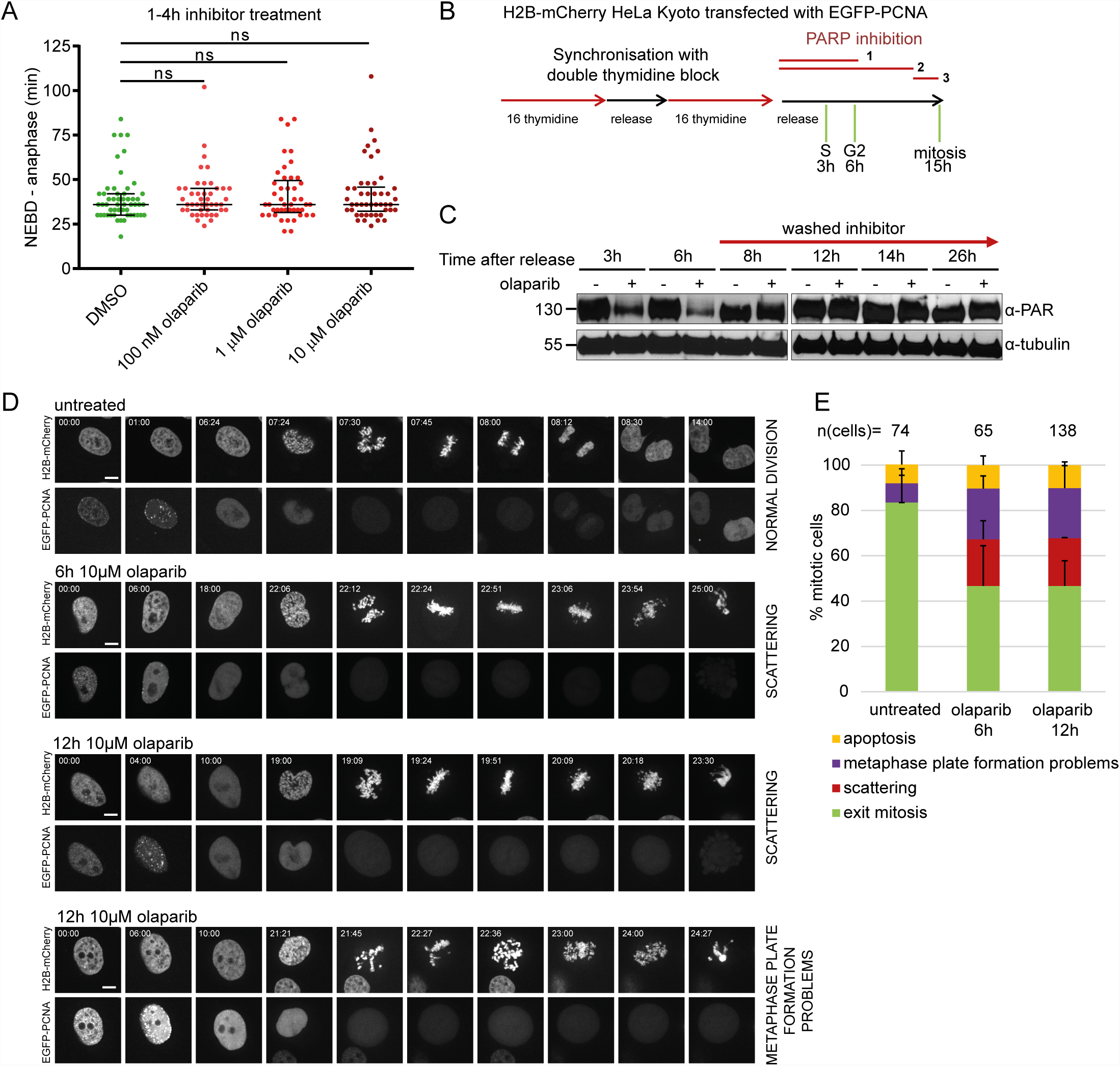
Premature loss of cohesion caused by olaparib is due to PARP inhibition in S-phase. **(A)** Duration of NEBD-anaphase analysed by live-cell imaging in HeLa cells stably expressing H2BmCherry EGFP-securin treated for 1-4 h with indicated concentrations of olaparib. **(B)** Experimental workflow for olaparib exposure during different stages of the cell cycle. Cells were synchronized by double thymidine block and olaparib was added ‘1’ during S-phase, ‘2’ during S and G2 phase, ‘3’ just before NEBD. **(C)** Western blot analysis showing efficient removal of olaparib at the end of S-phase. **(D)** Stills from spinning disc imaging and **(E)** quantification of representative mitotic phenotypes. Scale bar=10 μm.

### Olaparib directly induces premature loss of cohesion

To test whether olaparib induces premature sister chromatid separation directly by weakening sister chromatid cohesion, we compared the timing of scattering after olaparib treatment to other treatments previously shown to induce chromatid scattering (Figure 5A). In addition to MG132 and siCdc20, partial RNAi depletion of sororin, which is required for cohesion establishment in S-phase, was also shown to induce chromatid scattering (Supplementary Figure 4) [43, 44]. Cells arrested in metaphase with MG132 or siCdc20 exhibited chromatid scattering only after 4 h; however, siSororin rapidly induced scattering (1.2 h after NEBD), while 10 μM olaparib exhibited intermediate kinetics (2.4 h after NEBD) (Figure 5A). This indicates that olaparib treatment may directly weaken chromosome cohesion.

**Figure 5:**
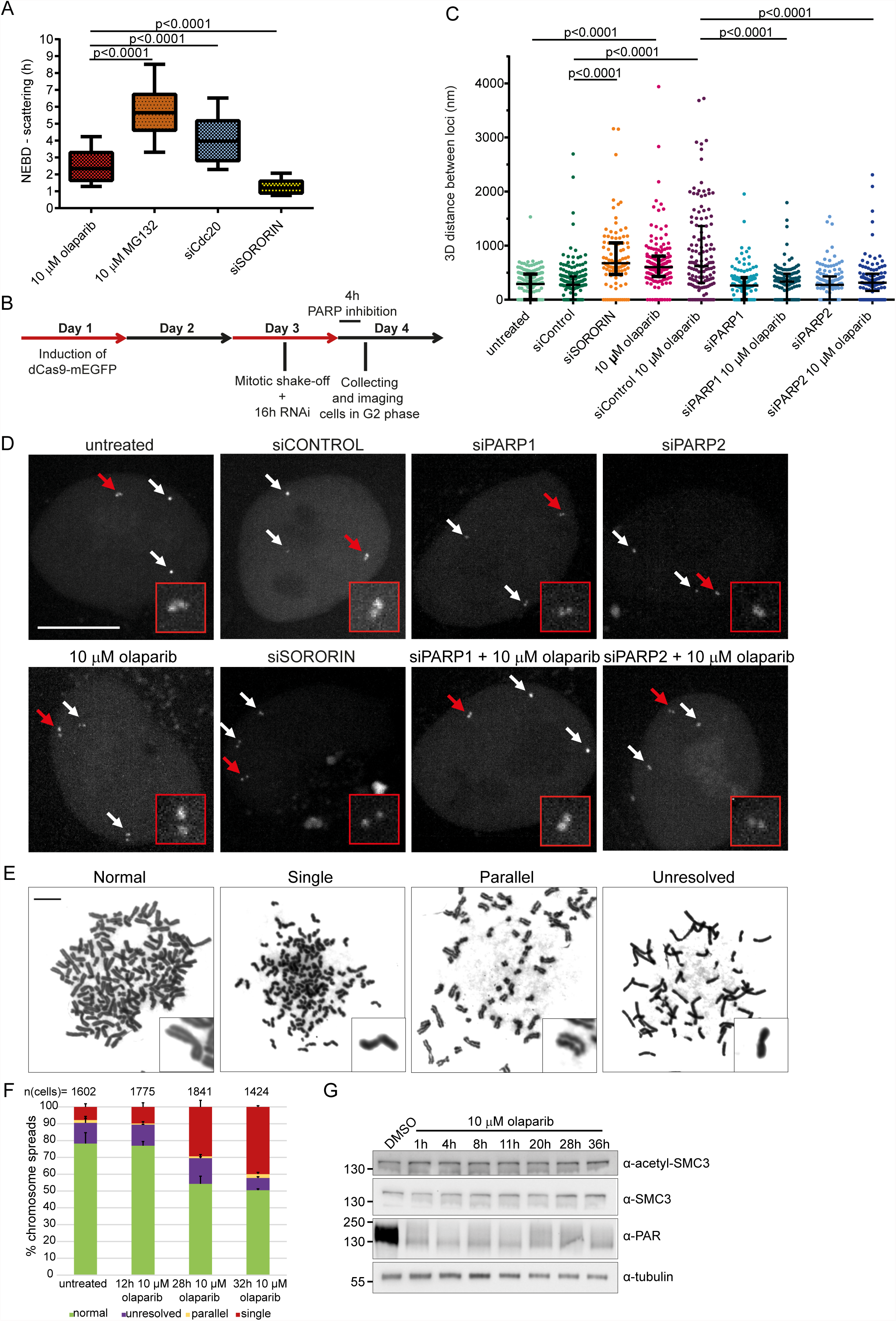
Scattering is a result of premature loss of cohesion caused by olaparib. **(A)** Comparison of NEBD-scattering duration induced by various treatments. Cells were treated with 10 μM olaparib for 24 h and 10 μM MG132 for 30 min. Cdc20 and sororin were depleted for 24 h with RNAi. Number of cells analysed per condition: n (olaparib) = 44, n (MG132) = 88, n (siCdc20) = 94, n (siSororin) = 63. **(B)** Experimental workflow for measuring sister chromatid distances. dCas9-mEGFP targeting Muc4 loci is induced, followed by mitotic shake-off, PARP inhibition for 4 h or RNAi for 16 h before imaging of G2 cells. **(C)** 3D distance between sister chromatids at Muc4 loci measured after live-cell imaging of PARP-inhibited or PARP/sororin-depleted cells. **(D)** Representative images quantified under **(C)**. Scale bar=10 μm. **(E)** Representative images and **(F)** analysis of phenotypes revealed by Giemsa-stained chromosome spreads after addition of 10 μM olaparib. Scale bar= 10 μm. **(G)** Olaparib does not affect SMC3 acetylation levels. Extracts from HeLa cells treated with 10 μM olaparib for various times were analysed by Western blotting.

To directly measure the effect of olaparib on sister chromatid distances during interphase, we visualized a genomic locus (Muc4) in live HeLa cells by stably expressing dCas9-mEGFP and guide RNAs targeting 3^rd^ exon of the Muc4 gene (hg19 Chr3: 195506180-195510888), as previously reported [45]. We synchronized cells by mitotic shake-off and imaged the labeled loci 16 h later, when cells were in G2 (Figure 5B). 64% of the labelled loci appeared as doublet dots (Figure 5C-D), consistent with a separation of the two replicated sister loci by a distance larger than the resolution limit of the confocal microscopy. The median distance between separated sister loci in control cells was 275 nm (n=195) and it increased in sororin-depleted cells to 670 nm (n=94) (Figure 5C-D). A 4 h-treatment of S-phase cells with 10 μM olaparib also substantially increased the distance between sister loci to 617 nm (n=136), whereas siPARP1 and siPARP2 had no effect (Figure 5B-D).

This confirms that S-phase PARP1/2 inhibition by olaparib directly weakens sister chromatid cohesion in interphase cells ultimately resulting in sister chromatid scattering in metaphase cells. Importantly, siPARP1 and siPARP2 rescued the effect of olaparib, confirming that olaparib-induced sister chromatid separation is caused by on-target inhibition of PARP1/2 (Figure 5B-D).

In addition, we analysed chromosome spreads from untreated and olaparib-treated HeLa cells collected by mitotic shake-off (Figure 5E-F). 7.7±1.9% of untreated spreads contained single chromatids, which likely represent anaphase stages (Figure 5E-F). 10 μM olaparib gave rise to 29.3±3.9% and 39.9±0.7% of spreads with single chromatids at 28 or 32 h after treatment, respectively (Figure 5E-F). This corresponds well to the percentage of cells exhibiting chromatid scattering (30±4% for 10 μM olaparib after 30-37 h) (Figure 1D).

Acetylation of SMC3 was shown to mediate cohesion establishment in S-phase by promoting sororin recruitment to the cohesin complex [43]. Thus we tested whether olaparib impairs cohesion establishment by examining SMC3 acetylation in olaparib-treated cells (Figure 5G). Acetylation of SMC3 did not change upon olaparib treatment, which suggests that olaparib does not impair cohesion establishment but instead causes loss of cohesion. Cohesion defects caused by depletion of spliceosome subunits did not affect SMC3 acetylation either, confirming that loss of cohesion can occur despite unperturbed cohesion establishment [46, 47].

Taken together, our data provide comprehensive evidence that the olaparib-induced scattering phenotype results from premature loss of sister chromatid cohesion in replicating cells.

### Olaparib causes chromatid scattering in several cancer cell lines

The above characterization of olaparib effects on mitotic progression was performed on HeLa cells as a commonly used cancer cell line of cervical origin. We extended our characterization to a set of non-cancerous cell lines (human mammary epithelial, HME1 [48]; retinal pigment epithelial, RPE1 [36]) and cells of cancerous origin from cervix (C33-A, SiHa), ovary (TOV-21G), breast (MDA-MB-468 [49] and BT-549) and osteosarcoma (U2OS [50]) (Supplementary Table S2) and visualized chromosomes in live cells with the non-toxic SiR-Hoechst DNA dye [51]. Anaphase delay and chromatid scattering in HeLa SiR-Hoechst-labelled cells treated with olaparib were comparable to HeLa H2B-mCherry cells (Supplementary Figure 5), validating that SiR-Hoechst is not toxic in this assay.

Given that inhibition of drug efflux channels by verapamil was previously shown to decrease the survival of MRN-deficient colon cancer cells (HCT116 cells) treated with a PARP inhibitor [52], we also tested whether verapamil would potentiate the mitotic phenotypes caused by olaparib in different cell lines (Figure 6).

**Figure 6:**
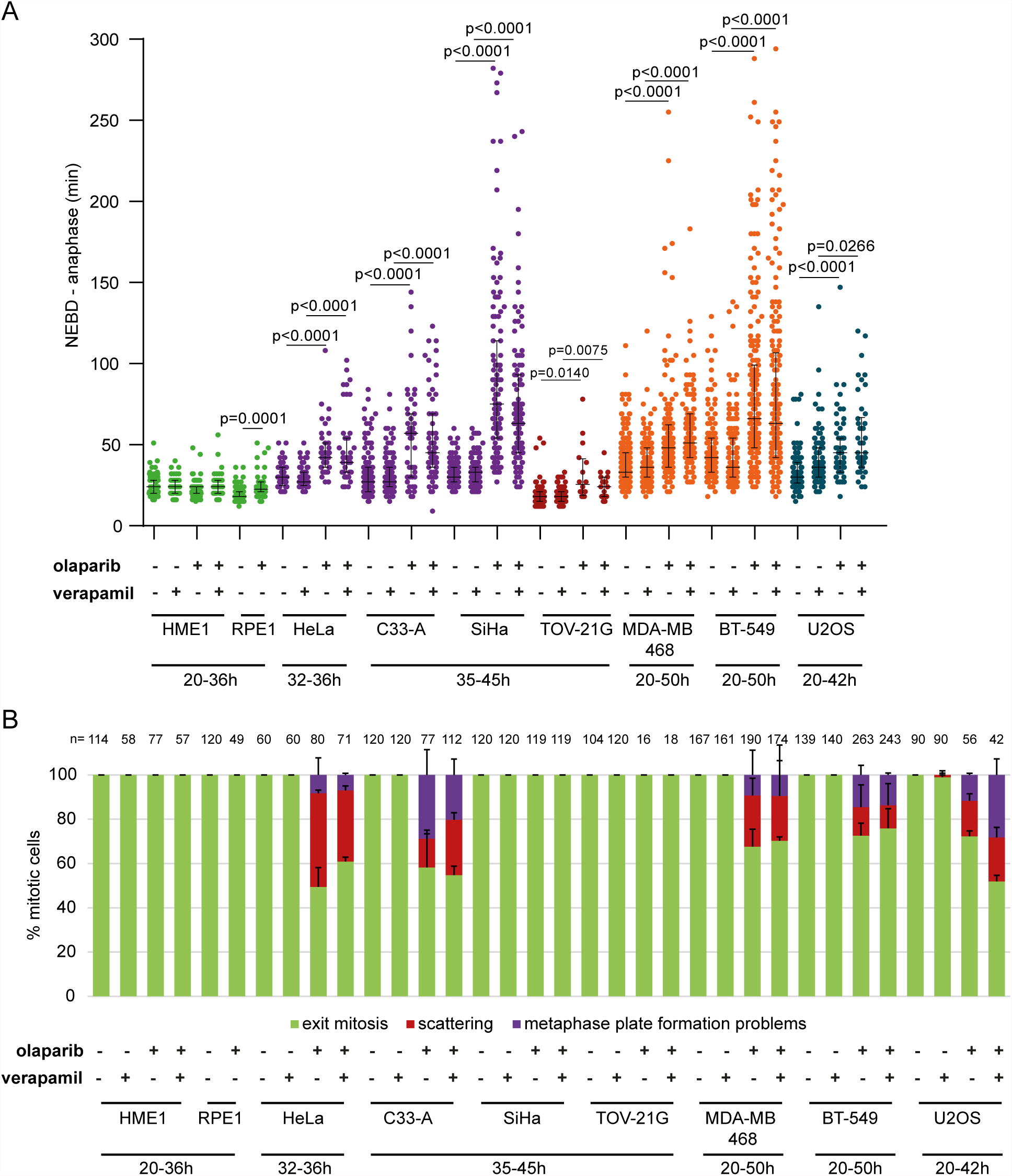
Olaparib causes anaphase delay and chromosome scattering in different cancer cell lines. **(A)** Duration of NEBD-anaphase in non-cancerous and cancerous cell lines after verapamil, olaparib or concomitant verapamil and olaparib treatment. Later time points were analysed in cell lines that were highly stalled by olaparib treatment. Green = non-cancerous cell lines, violet = cervical cancer cell lines, dark red = ovarian cancer cell line, orange = breast cancer cell lines and blue = osteosarcoma cancer cell line **(B)** Percentage of the observed mitotic phenotypes.

Olaparib reduced PARP1 auto-PARylation and total PARylation in all tested cell lines (Supplementary Figure 6). 10 μM olaparib did not induce scattering in HME1 GFP-H2B and RPE1 mRFP-mCherry cells, while RPE1 cells exhibited a minor anaphase delay (7 min; Figure 6). Olaparib treatment in TOV-21G resulted in anaphase delay (45 min and 8 min, respectively) without causing chromatid scattering (Figure 6). Anaphase delay coupled with chromatid scattering was observed in C33-A, MDA-MB-468, BT549 and U2OS cells (Figure 6). Chromatid scattering was most pronounced in HeLa (32±2%), followed by C33-A (25±3%), MDA-MB-468 (20±16%), U2OS (20±5%), and BT-549 (11±10%) after concomitant treatment with olaparib and verapamil (Figure 6B).

Chromatid scattering does not correlate with the p53 status as HeLa and U2OS cells, which showed scattering, have a wild-type p53 (Supplementary Table S2 and Supplementary Figure 7). To examine whether the degree of olaparib-induced chromatid scattering correlates with its cytotoxicity, we determined survival of all cell lines after exposure to a range of olaparib concentrations using colony formation assay and MTS assay to determine 14-day and 3-day cytotoxicity, respectively (Supplementary Figure 8). The degree of scattering correlated with cytotoxicity and HeLa was most sensitive to olaparib (Supplementary Figure 8). However, cell lines that did not show scattering or other mitotic phenotypes, such as RPE1 and TOV-21G, were more sensitive to olaparib than HeLa, which suggests that in these cell lines olaparib induces cell death via non-mitotic pathways (Supplementary Figure 8). This is not surprising given that different agents can cause cell death via different pathways (e.g., apoptosis, necrosis, mitotic catastrophe, autophagy) in different cell types depending on their genotypic and phenotypic properties [53, 54].

### Chromatid scattering correlates with high PARP1 and PARP2 expression levels

The degree of cytotoxicity of PARP1 inhibitors was previously found to correlate with PARP1 expression levels, which are known to be increased in breast cancer [55]. Reduced PARP1 expression levels were associated with resistance to PARP inhibitors [56]. To test whether the scattering phenotype correlates with PARP expression levels, we quantified PARP1 and PARP2 mRNA levels in different cell lines by RT-qPCR and PARP1 and PARP2 protein levels by Western blotting (Figure 7A,B). Both mRNA and protein levels were very high in cell lines of cervical origin, particularly HeLa and C33-A, which also exhibited the highest degree of chromatid scattering (Figure 7A,B). Cell cycle analysis by flow cytometry revealed olaparib-induced S-phase stalling and G2/M arrest in all tested cell lines with the exception of HME1 (Figure 7C). Collectively, these data show that olaparib-induced chromatid scattering during mitosis correlates with high PARP1 and PARP2 expression levels (Figure 7D).

**Figure 7:**
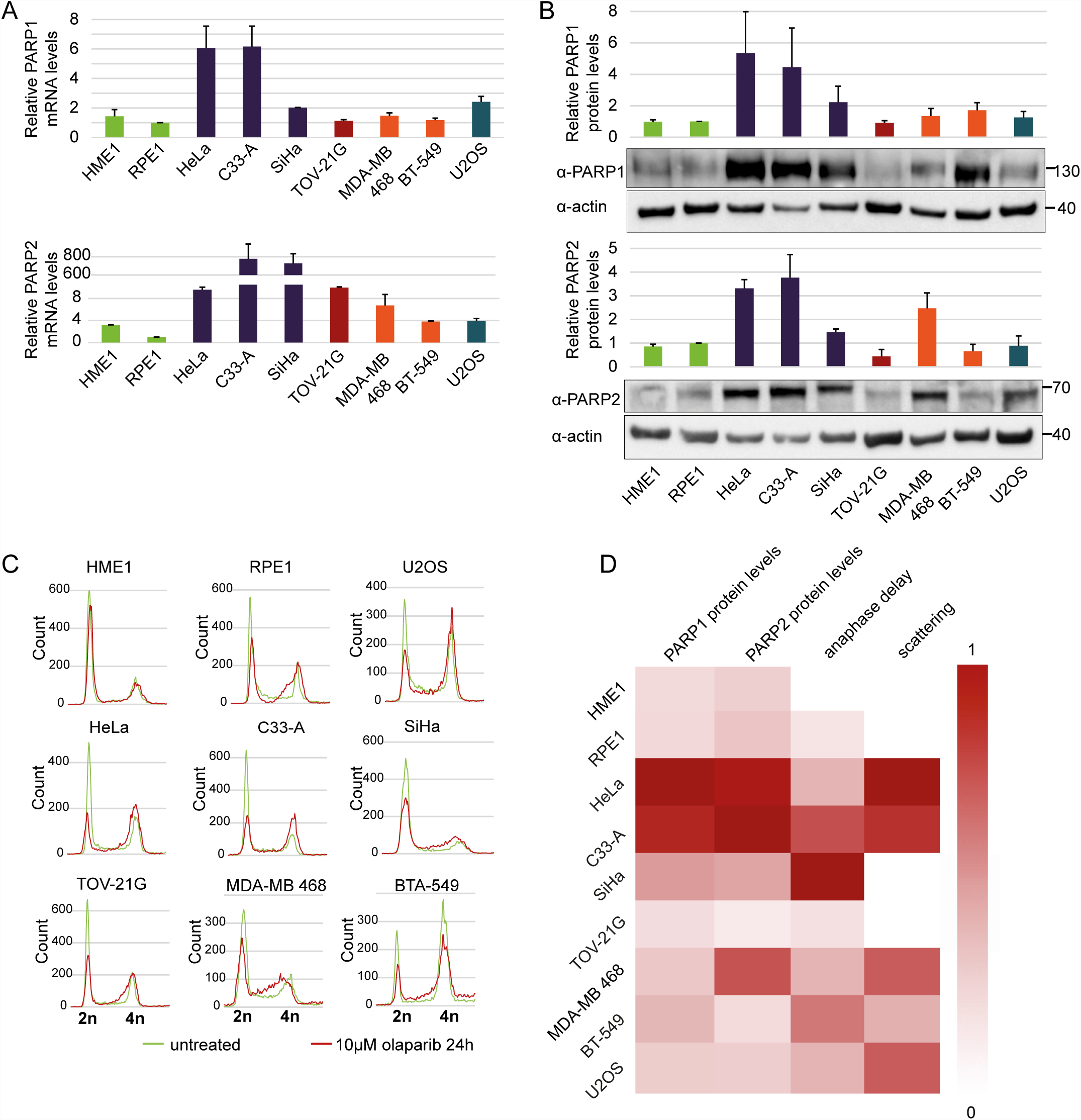
Cervical cancer cell lines HeLa and C33-A have high PARP1 and PARP2 mRNA and protein levels and are arrested in G2/M phase after olaparib treatment. **(A)** Relative PARP1 and PARP2 mRNA levels normalized to actin as a loading control. **(B)** Western blot analysis of PARP1 and PARP2 protein levels normalized to actin as a loading control. **(C)** Flow cytometry analysis of cancerous and non-cancerous cell lines after olaparib treatment. **(D)** Heat map showing correlations between anaphase delay and scattering caused by olaparib and relative PARP1 and PARP2 protein levels. All values were normalized from 0-1. Dark red (1) represents the highest protein levels, strongest anaphase delay and highest degree of scattering, while white (0) represents the lowest values.

### Chromatid scattering does not correlate with the degree of DNA damage caused by olaparib

Given that cancer cells may have increased proliferation and increased levels of replication stress, which makes them more sensitive to ATR or Chk1 inhibitors [57, 58], we tested whether the propensity of different cell lines towards chromatid scattering correlates with elevated growth rate and elevated replication stress upon olaparib treatment. The growth rate was determined by counting the number of cells over four days (Figure 8A). We did not observe a correlation between the proliferation rate and the degree of scattering upon olaparib treatment among different cell lines (Figure 8A). To determine the levels of replication stress, we measured the total levels of phosphorylated RPA (replication protein A), which is found at single-stranded DNA generated at replication forks upon uncoupling of DNA helicase from DNA polymerase [59]. Phospho-RPA levels were two-fold increased in olaparib-treated HeLa and C33-A, which also showed the highest degree of chromatid scattering (Figure 8B and Figure 6). Other cell lines did not show a correlation between phospho-RPA and scattering (Figure 8B). We also compared the number of γH2AX foci-positive cells in untreated versus olaparib-treated cells and observed a similar increase in all cell lines (Figure 8C), as previously reported for olaparib-treated malignant lymphocyte cell lines [40]. γH2AX foci were also found in olaparib-treated mitotic cells that exhibited chromatid scattering (Figure 8D). However, the levels of γH2AX foci did not correlate with the degree of scattering upon olaparib treatment (Figure 8C). Differences in sensitivity to olaparib-induced chromatid scattering can thus be partially linked with the induction of replication stress but not the proliferation rate nor the levels of DNA damage. Replication stress was recently also detected upon olaparib treatment of MYCN-amplified neuroblastoma, which also express high PARP1/2 levels [60]. This provides further support for the model whereby high PARP1 and PARP2 levels are conducive to PARP entrapment and replication blockage resulting in chromatid scattering after olaparib treatment.

**Figure 8:**
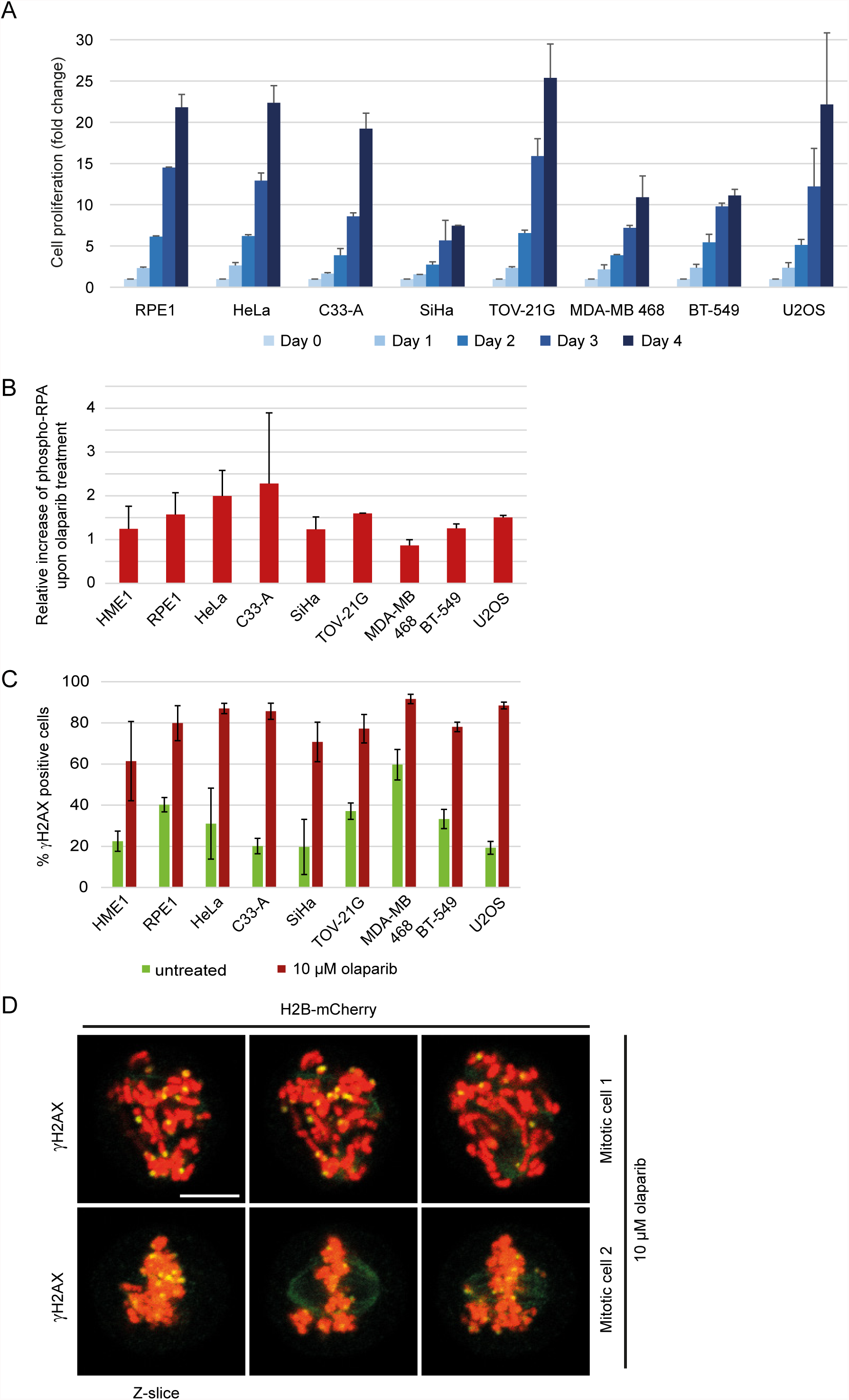
Replication stress and DNA damage after olaparib treatment. **(A)** Growth rate of various cell lines determined by cell counting over 4 days. **(B)** Relative increase of phospho-RPA levels in untreated vs 10 μM olaparib-treated cells for 30 h. Immunofluorescent phospho-RPA intensities were measured by ImageJ (n>100). **(C)** Percentage of γH2AX positive cells in untreated and 10 μM olaparib-treated cells for 30 h. Cells were scored as γH2AX positive if the number of foci per nucleus was ≥5 (n>100). **(D)** Representative images of γH2AX foci in mitotic cells treated with 10 μM olaparib for 30 h. Scale bar=10 μm.

### Sister chromatid scattering is a common outcome of replication fork perturbation

In order to assess whether chromatid scattering induced by PARP1/2 inhibitors is a common consequence of replication stress-inducing agents, we tested the effect of hydroxyurea, cisplatin, campthotecin and etoposide on cell division. Hydroxyurea induces replication fork stalling by inhibiting ribonucleotide reductase and thereby depleting dNTP pools [61]. Cisplatin generates DNA crosslinks, while camptothecin and etoposide inhibit topoisomerase I and II thereby inducing singlestrand and double-strand DNA breaks, respectively [62, 63]. Unlike hydroxyurea, cisplatin, camptothecin and etoposide directly damage DNA, which causes replication fork stalling and collapse in dividing cells [62-64]. To test the effect of these agents on mitotic progression by live imaging, we used the lowest concentration that induced pronounced S-phase stalling according to flow cytometry (Supplementary Figure 9). All agents induced chromatid scattering and metaphase plate formation problems, albeit to a different extent; hydroxyurea resulted in comparable levels of chromatid scattering and metaphase plate formation problems, whereas metaphase plate formation problems was the predominant mitotic phenotype of cisplatin, camptothecin and etoposide (Figure 9A-B). This suggests that chromatid scattering is a common outcome of replication stress-inducing agents, whereas metaphase plate formation problems arises as a dominant phenotype of DNA-damaging agents.

**Figure 9:**
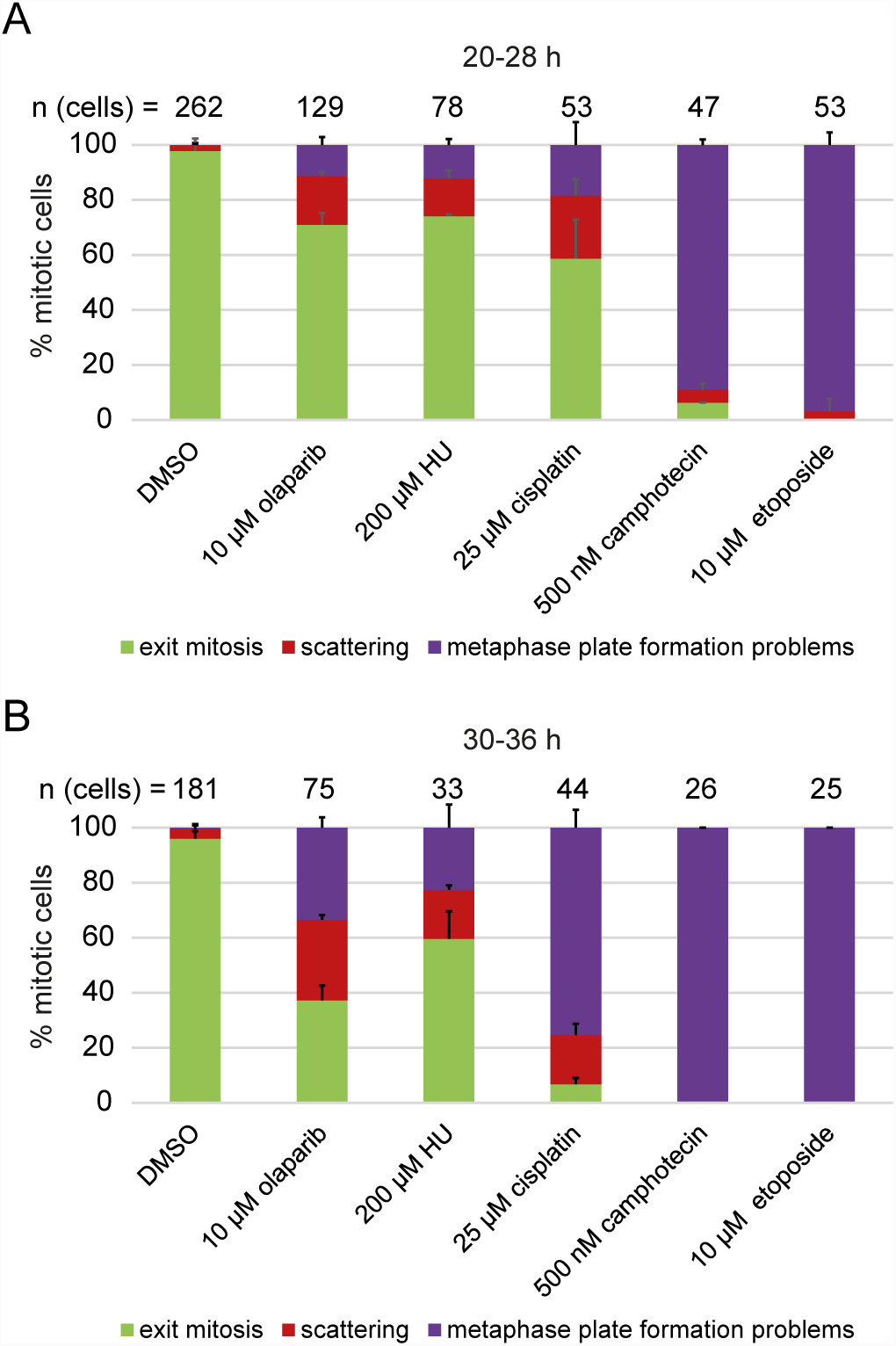
Sister chromatid scattering is a general outcome of replication fork perturbations induced by various agents. **(A,B)** Percentage of different mitotic phenotypes observed after **(A)** 20- 28 h or **(B)** 30-36 h treatment with replication stress-inducing agents.

## MATERIALS AND METHODS

### RNAi and inhibitors

Cells were seeded and transfected 24 h prior to live imaging in the presence of different siRNAs (30-50 nM) using RNAiMAX (Invitrogen). EGFP-PCNA transfection was performed 24 h prior to imaging using FugeneHD (Promega). The following inhibitors were used: olaparib (AZD2281, Ku-0059436; Selleckchem), veliparib (ABT-888; Selleckchem), talazoparib (BMN 673; Selleckchem); XAV-939 (Selleckchem), ME328(Lindgren et al 2013), verapamil (Sigma), MG132 (Sigma), hydroxyurea (Sigma), cisplatin (CPDD, Sigma), (S)-(+)-camptothecin (Sigma), etoposide (Sigma). SiR-Hoechst (Life Technologies) was used at the final concentration of 500 nM.

### Generation of EGFP-PARP1 and EGFP-PARP2 overexpressing cell lines using lentiviral system

EGFP-PARP1 and EGFP-PARP2 were cloned into a transfer plasmid under the EF1a promoter. HEK 293NT cells were transfected with transfer plasmid, viral envelope coding plasmid and lentiviral packaging plasmid using PEI. 36 h after transfection virus-containing supernatant was filtered and added to HeLa RIEP cells (Day 1). Second infection was performed 24 h later (Day 2). On Day 3 cells were washed with PBS and placed in fresh media. On Day 6 washing was repeated and GFP-positive cells were FACS sorted.

### Microscopy

Live-cell wide-field microscopy experiments on Molecular Devices ImageXpress Micro screening microscope were performed using 96-well plates as already described [51]. Live-cell spinning disc microscopy was performed using 4-well glass bottom dishes (Greiner) in an environmentally controlled chamber with an Axio Observer Z1 (Zeiss) inverted microscope equipped with an EM-CCD camera (Evolve EM-512), Yokogawa CSU-X1-A1 spinning disc unit (pinhole diameter 50 μm), 488 nm diode laser, 561 nm DPSS laser (AOTF-controlled) and a Plan-Apochromat 63x/1.4 oil-immersion objective. Cells were maintained at 5% CO2 and 37°C during experiments. Confocal microscopy was performed on a customized Zeiss LSM 710 microscope using a x63, 1.4N.A oil Plan-Apochromat objective (Zeiss).

### Immunofluorescence

Cells were grown on high-precision borosilicate cover glasses (LH22.1 Roth Labware). For cyclin B staining (1:200; Cell Signalling), cells were fixed in PTEMF buffer or in 4% PFA followed by ice-cold methanol, and stained as previously described [29]. For γH2AX and phospho-RPA staining, cells were fixed in 4% PFA, permeabilised in 0.5% Triton X-100 in PBS and blocked for one hour in 0.1% Tween and 1% BSA, followed by incubation with primary antibodies: rabbit anti-γH2AX (1:500; Bethyl), rabbit anti-phospho-RPA S4/S8 (1:500; Bethyl). and subsequent incubation with appropriate secondary antibodies (1:500; Alexa Fluor^®^).

### Image and statistical analysis

Image analysis was performed using FiJi (ImageJ v1.5). Duration from NEBD (nuclear envelope breakdown) to anaphase (min) is presented using median with interquartile range. In Figure 5a box and whiskers are plotted and presented as 10-90 percentile. Statistical analysis was performed using Kolmogorov-Smirnov test. Live imaging microscopy data are based on at least two biological replicates with three or more technical replicates each.

### FACS

Cell cycle profiling was performed using propidium iodide staining as described[46], measured at Zytofluorometer FACSCalibur and analysed using Flowing Software 2.5.1.

### Sister chromatid distance measurements

5x10^5^ Muc4 TRE3G-dCas9-mEGFP cells [45] were seeded in T75 flasks and dCas9-mEGFP expression induced with 1 μg/mL doxycycline for 48 h. Shake-off and collection of mitotic cells was performed as follows. After a pre-shake-off and 2x wash with pre-warmed PBS, the cells were allowed to enter mitosis for 2 h in pre-warmed medium with 1 μg/mL doxycycline. Mitotic cells were mechanically detached by shaking the plates, centrifuged at 500 g for 5 min, resuspended in an appropriate amount of medium containing 1 μg/mL doxycycline and seeded in 4-chamber glass bottom dishes (Greiner). For silencing experiments siRNA and RNAiMAX reagent were mixed and incubated as described above and added to the glass bottom dish prior to seeding mitotic cells. Live imaging was performed 16 h after the mitotic shake-off when the cells entered G2 phase. 10 μM olaparib was added 3 h prior to imaging when the cells were in S-Phase (13 h after mitotic shake-off). Imaging was performed on a spinning disc microscope with an sCMOS 2xpco.edge 4.2 camera (0.065 μm pixel size) and an EC Plan-Neofluar 100x/1.3 oil-immersion objective. Z-stack images were acquired in 100 nm intervals. The 3D distance of paired sister chromatids was determined with the Fiji Plugin Trackmate (DoG detector, sub-pixel localization, estimated spot diameter of 150 nm) by using the x, y and z coordinates of the detected sister chromatids to calculate the distance d=sqrt((X_1_-X_2_)^2^ + (Y_1_-Y_2_)^2^ + (Z_1_-Z_2_)^2^).

### Chromosome spreads

Chromosome spreading and Giemsa staining was performed as previously described[46, 65], excluding the nocodazole treatment. In total over 6600 prometaphase/metaphase spreads were analyzed in two independent experiments (each containing five technical replicates) by automated analysis using Metafer4 v 2.12.116 (MetaSystems). All slides were scanned automatically and spreads were randomly selected in each technical replicate using Metafer4. Spreads were then blindly categorized into four phenotypes.

### Quantitative RT-PCR

Total RNA was extracted with TRI reagent (Sigma) and PCl (Phenol equilibrated, stabilized: Chloroform: Isoamyl alcohol 25:24:1; AppliChem), and precipitated in ethanol. RT (reverse-transcription) reaction was performed with random primers (Invitrogen 48190_011) and 5xProtoScript II RT (BioLabs). Quantitative PCR was performed with 5xHot FirePol Eva Green qPCR mix (BioZyme). All experiments were repeated twice with two technical replicates.

### Immunoblotting

Cell pellets were lysed in 50 mM Tris pH8, 150 mM NaCl, 1% Triton, 1 mM DTT, 50 units/ml benzonase (Novagen) and protease inhibitors (EDTA-free, Roche). Mouse anti-tubulin (1:5000; Sigma), mouse anti-actin (1:20000; Sigma), rabbit anti-PARP1 (1:1000; Cell Signalling), rabbit anti-PAR (1:1000; Trevigen), mouse anti-PARP2 (1:50; Enzo), rabbit anti-PARP3 (1:200; Dantzer lab), rabbit tankyrase-1/2 H-350 (1:200; Santa Cruz), rabbit anti-Cdc20 (1:1000; Peters lab), rabbit anti-human sororin (1:1000; Peters lab), rabbit anti-SMC3 (1:1000; Bethyl laboratories), mouse anti-acetyl-SMC3 (1:1000; Shirahige lab), rabbit anti-P53 (1:1000; Cell Signaling), rabbit anti-BRCA1 (1:1000; Cell Signaling), rabbit anti-BRCA2 (1:1000; Cell Signaling). Secondary HRPconjugated antibodies (Jackson ImmunoResearch) or IRDye fluprescent dye antibodies (LI-COR) were used at 1:10000 dilution. Images were taken on ChemiDoc Touch Imaging System (Bio-Rad) or Odyssey CLx imager (LI-COR) using Image Lab 5.2.1 for analysis.

### Colony formation assay

Cells were seeded and treated with different concentrations of olaparib at a very low confluence (1000-3000 cells per 60 mm dish). Medium (with or without olaparib) was exchanged every 4-5 days. After 14 days, medium was removed and cells were fixed with 4% PFA and incubated 10 min RT. PFA was removed and 0.1% crystal violet in 25% methanol was added to the dish and incubated 20 min at 4°C. Crystal violet was removed and dishes were washed in filtered water until residual crystal violet was completely removed. Percentage of surviving cells was quantified by measuring the area of colonies using the ImageJ-plugin ‘ColonyArea’. The intensity of colonies was not taken into account due to variable staining of different cell lines by crystal violet.

### MTS assay

HeLa cells were seeded in 96-well plates (1000 cells per well) in the presence of siRNAs. Olaparib was added 24 h later. For the comparison of survival across cell lines, RPE1, HME1, HeLa, TOV-21G, BT-549 and U2OS cells were seeded at 1000 cells per well; SiHa was seeded at 2000 cells per well; MDA-MB-468 was seeded at 4000 cells per well; C33-A was not measured as it does not metabolize the MTS reagent. Olaparib was added during seeding. CellTiter 96^®^ solution (Promega) was added after 72 h and the absorbance was measured at 490 nm.

### Proliferation assay

5x10^4^ cells were seeded in one well of a 6-well plate on day zero. After 24 h cells were trypsinized, resuspended in DMEM media and counted using Mini Automated Cell Counter (ORFLO). Counting was repeated each 24 h for 4 consecutive days. Cell proliferation was quantified by normalizing the number of cells with the number of cells on day zero.

## DISCUSSION

Numerous clinical trials have been conducted since the initial development of PARP inhibitors [66], which culminated in the approval of olaparib for the treatment of BRCA-mutated ovarian cancer [67]. However, the molecular basis of PARP inhibitor function remains unclear [68, 69]. PARP1, PARP2, PARP3 and PARP5a (tankyrase) co-localization with various mitotic structures prompted us to study the effect of their inhibition or depletion on mitosis. By tracking mitotic progression of individual cells, we uncovered a new phenotype of PARP1/2 inhibition by olaparib. We showed that, by acting on replicating cells, olaparib induces metaphase arrest and sister chromatid scattering, ultimately resulting in cell death (Figure 10). Moreover, we delineated the mechanism of mitotic scattering by showing that olaparib causes loss of sister chromatid cohesion in G2 cells. The olaparib-induced scattering phenotype is suppressed by PARP1 or PARP2 depletion, demonstrating that PARP1 and PARP2 are the relevant targets and that PARP1/2 trapping rather than catalytic inhibition is the mechanism of olaparib-induced mitotic failure. Chromatid scattering was observed in various cancer cell lines and was correlated with PARP1 and PARP2 expression levels.

**Figure 10:**
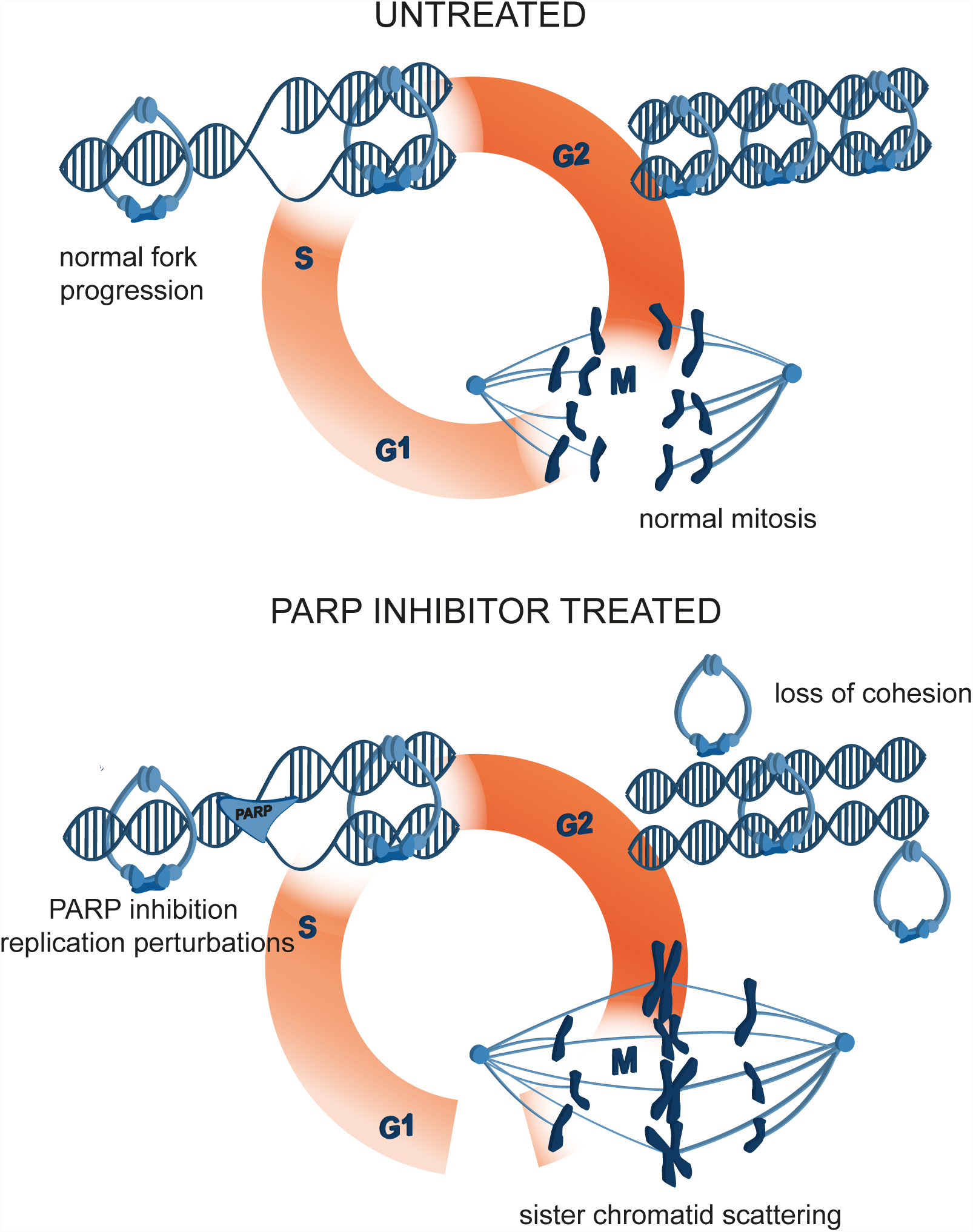
A model of mitotic cell death caused by premature loss of cohesion due to PARP inhibition with olaparib. PARP inhibition causes replication fork stalling and premature loss of cohesion in interphase. Mitotic cells are consequently arrested in metaphase, sister chromatids scatter away from the metaphase plate and the cells eventually die.

The PARP inhibitor olaparib is the first clinically approved drug that targets DNA damage response (DDR) [67]. Olaparib is particularly efficient in tumors lacking specific DDR functions, which renders them more reliant on a particular DDR pathway. In particular, patients bearing mutations in BRCA1/BRCA2 genes that are required for the homologous recombination DNA repair pathway have shown exceptional susceptibility to PARP inhibitors. Multiple clinical trials carried out since 2009 have demonstrated the beneficial effects of olaparib on BRCA-mutated ovarian and breast cancer, as well as prostate and pancreatic cancer, Ewing’s sarcoma, small cell lung carcinoma and neuroblastoma [18, 70]. Given that cancers without mutations in DNA repair pathways are also susceptible to PARP inhibition, non-DNA repair functions of PARP1/2 are likely also responsible for the deleterious effects of PARP inhibition [18]. Indeed, our study shows that olaparib induces chromatid scattering in metaphase-arrested cells resulting in cell death. However, the observed olaparib-induced mitotic phenotype is not caused by PARP1/2 inhibition in mitosis but instead results from PARP1/2 inhibition during S-phase replication.

Contrary to olaparib treatment, PARP1 or PARP2 depletion did not result in chromatid scattering. Differential effects of PARP inhibition and PARP depletion have been attributed to the entrapment of PARP1 on DNA by inhibitors such as olaparib and talazoparib [13]. By interacting with the D-loop at the outer border of the NAD site, olaparib and talazoparib induce conformational changes in the PARP1 DNA-binding domains to stabilize the PARP1-DNA complex [71]. Trapped PARP-DNA complexes prevent DNA replication and transcription; PARP poisoning effect therefore determines the potency of PARP inhibitors rather than their effect on PARP catalytic inhibition [13, 68]. As chromatid scattering results from olaparib-induced loss of cohesion in S/G2 phase that can be rescued by PARP1/2 depletion, we surmise that olaparib-induced chromosome scattering is caused by PARP1/2 trapping. This is further supported by a correlation between high PARP1 and PARP2 levels, induction of replication stress and chromatid scattering in HeLa and C-33A cell lines.

Replication problems have been already linked with mitotic defects [72]. Faithful segregation of sister chromatids can be compromised by incompletely replicated chromosomes caused by replication fork stalling, incompletely resolved DNA repair intermediates or topologically intertwined sister chromatids [72]. Mitotic structures caused by replication problems include anaphase chromatin bridges and ultrafine DNA bridges [72]. So far, metaphase arrest and premature loss of cohesion have not been linked with replication problems. By extending our analyses to other agents that perturb replication fork progression, such as hydroxyurea, cisplatin and topoisomerase inhibitors, we showed for the first time that replication problems can also lead to premature loss of cohesion in metaphase-arrested cells.

Genetic predisposition (e.g., BRCA mutations) or phenotypic characteristics (e.g., platinum resistance) are not sufficient to predict patient response to olaparib treatment [58]. Chromatid scattering was consistently observed in cervical cancer cells with increased PARP1 and PARP2 protein levels (HeLa, C33-A), suggesting that PARP1 and PARP2 protein levels could be used as a predictive biomarker for the efficiency of olaparib treatment, as previously proposed [60, 68]. Olaparib is currently undergoing various clinical trials, including different gynaecological malignancies such as cervical and uterine cancer [73], where such a biomarker may be particularly useful.

In summary, we showed that sister chromatid scattering in mitosis is a new mechanism of olaparib-induced cytotoxicity. By entrapping PARP1 on replicating DNA, olaparib obstructs replication fork progression resulting in loss of sister chromatid cohesion in G2 cells (Figure 10). Loss of cohesion in interphase cells causes chromatid scattering in metaphase cells, metaphase arrest and cell death.

## AUTHOR CONTRIBUTIONS

E.K. designed, performed and analysed experiments and wrote the manuscript. T.K. performed spinning disc microscopy, γH2AX quantification, sister chromatid distance measurements and a part of live imaging screens. A.E.D. performed the initial live imaging experiment with olaparib. R.Z. provided the following cell lines: C33-A, SiHa, TOV-21G, BT-549. D.W.G designed live-cell imaging experiments and wrote the manuscript. D.S. designed and supervised experiments and wrote the manuscript.

## ACKNOWLEDGEMENTS

We thank Jan-Michael Peters, Miroslav Ivanov Penchev and Rene Ladurner for sharing reagents and helpful discussions. We thank Claudia Blaukopf for generating PARP1- and PARP2-overexpressing HeLa. We thank Rudolf Drescher (MetaSystems) for 3.12.116v 4Metafer; Sebastien Herbert for help with analysis of sister chromatid distances; Herwig Schüler for sharing ME328 inhibitor; Françoise Dantzer for PARP3 antibody; Ben Bouchet, Arshad Desai, Stephan Geley and Jean Zhao for sharing cell lines.

